# Hormone-Dependent Microstates and the Reconfiguration of Resting-State Dynamics across the Menstrual Cycle

**DOI:** 10.64898/2026.03.06.707993

**Authors:** Matteo Demuru, Marianna Angiolelli, Emahnuel Troisi Lopez, Mario De Luca, Enrica Gallo, Charlotte Maschke, Damien Depannemaecker, Laura Sarno, Carmine Granata, Pierpaolo Sorrentino

## Abstract

While behavioral fluctuations across the menstrual cycle (MC) are well-documented, the neural underpinnings of these changes remain elusive. This study investigated the hypothesis that cyclic variations in sex hormones modulate large-scale brain activation patterns. To test this, longitudinal magnetoencephalographic (MEG) recordings were acquired and source reconstructed from 24 naturally cycling women across three distinct MC phases: early follicular, peri-ovulatory, and mid-luteal. Microstate analysis was employed to characterize large-scale cortical dynamics as “visits” to specific global configurations (i.e., maps) of brain activity. Our results revealed significant variations in the occurrence of specific microstate maps, particularly between the early follicular and mid-luteal phases. Furthermore, the occurrence of these specific configurations was significantly associated with fluctuations in hormone levels. Critically, both the hormonal levels and microstate dynamics were predictive of individual longitudinal changes in psychological well-being. These findings propose a neurophysiological substrate for the behavioral effects of hormonal cycling, identifying specific topographic maps whose dynamics are sensitive to the hormonal profile and carry predictive power for psychological health. Collectively, these results underscore the necessity of accounting for the MC in neuroimaging research and introduce a novel framework for defining microstates (Hormone-Dependent Microstates - HDMs) with respect to slowly changing dynamical properties across a month-long timescale.

## Introduction

Fluctuations in mood and cognition are common features of the natural menstrual cycle (MC) in a substantial portion of the population^1–3^. These fluctuations range from subtle changes to overt disorders, such as Premenstrual Dysphoric Disorder (PMDD), where women experience severe, debilitating affective and cognitive symptoms during the luteal phase^4^.

Broadly, the effects of sex hormones can be categorized as either organizational or activational^5^. Organizational effects are typically long-term, developmentally driven, and constant, influencing both brain and behavior. In contrast, activational effects are transient and occur in response to fluctuations in hormone concentrations. These activational effects are particularly relevant to the MC, during which hormones such as follicle-stimulating hormone (FSH), estradiol (E), luteinizing hormone (LH), and progesterone (P) regulate the progression through the menstrual, follicular, ovulatory, and luteal phases. However, how these hormonal fluctuations reverberate across the brain to generate behavioural changes remains largely unknown.

To date, diverse neuroimaging modalities have been employed to characterize these effects, given the evidence that changes in large-scale dynamics are linked to changes cognitive and humoral changes. While research has primarily utilized functional and structural magnetic resonance imaging (MRI) to map hemodynamic and anatomical changes^2^, there is a growing shift toward electrophysiological techniques, such as electroencephalography (EEG)^1^ and magnetoencephalography (MEG)^6–12^. These modalities offer distinct perspectives on neural modulation. While fMRI-based approaches rely on indirect measures of brain activities via the hemodynamic response function, electrophysiological devices like EEG/MEG provide a direct window into synaptic activity^13^.

Multiple frameworks can be deployed to characterize these dynamics. Traditional analyses often rely on coarse temporal resolutions and use windowing techniques that assume stationarity (i.e., power spectral properties or functional connectivity)^14^. However, converging evidence has demonstrated that the brain is far from stationary and, rather, its dynamics are multistable, that is, they evolve over multiple stationary states^15,16^. As such, it might be appropriate to utilize approaches that do not aggregate over time but instead capture the trajectories of the states visited. In particular, brain rhythms have been shown to evolve over multiple timescales, with faster dynamics nested within slower activities, ranging from milliseconds to potentially years^17–19^.

The MC has been shown to influence brain connectivity, dynamics, and structure, with recent studies highlighting the role of sex-hormone fluctuations in modulating neural activity. For instance, in a longitudinal study, fluctuations of estradiol and progesterone have been linked to changes in white matter microstructure, cortical thickness, and tissue volumes, suggesting structural plasticity across the MC^20^. fMRI connectivity findings, however, remain mixed, with some studies reporting stability in resting-state networks^21–24^, while others demonstrate hormone-driven changes in the default mode, executive control, and auditory networks^25–28^. EEG and MEG studies further reveal dynamic reorganization of large-scale brain networks^10^, with MC phases influencing power spectra^6,8,9,29^, hemispheric asymmetry^30^, and effective brain connectivity^31^.

In the present study, we aim to investigate the impact of the MC on neural dynamics in naturally cycling women, assessed using source MEG-based microstate analysis to capture the sub-second transitions of quasi-stable brain states^32^. While previous electrophysiological research has focused predominantly on time-averaged spectral characteristics^1^, this framework detects transient fluctuations in neural architecture^11,33^. Critically, we address a significant gap^1,2^ in the field by coupling these neural measures computed across the MC with behavioral assessments.

We employed source-level microstate analysis of longitudinally recorded data across three phases of the MC—early follicular, peri-ovulatory, and mid-luteal—to examine whether specific functional topographical brain configurations (microstates) are uniquely associated with these phases.

Inspired by the methodology developed by Tait et al. (2022)^32^ the primary goal of our study was not to describe the four to five canonical brain activity clusters^34^, but rather to identify and characterize specific states that are maximally responsive to the phase of the MC. Therefore, we first extracted the Global Field Power peaks (GFP) and clustered them (where each cluster represents a state). Then, we moved away from the classical pipeline, and set out to identify microstates that could differentiate the early follicular, peri-ovulatory, and mid-luteal phases based on their total number of occurrences across subjects. To this end, we identified the optimal number of states so that most of the variations associated with the menstrual phase would collapse into a minimal number of states (the Hormone-Dependent Microstates - HDMs). Finally, we evaluated, at the single-subject level, whether the changes occurring along the MC in the number of visits to the HDMs is predictive of the behavioral outcomes, as assessed by six dimensions of well-being (autonomy, environmental mastery, personal growth, positive relations with others, purpose in life, and self-acceptance).

## Results

### Hormones across the menstrual cycle phases

The variations in blood hormone concentrations (LH, FSH, P, and E) across menstrual cycle phases have been previously reported in Liparoti et al. (2021)^10^, we provided additional descriptive statistics in the supplementary materials Table S1.

As a sanity check, we showed that the hormonal profile change throughout the MC (*F*(2, 46) = 212.4, *p* < 0.001), in all phase pairs: peri-ovulatory vs. early follicular (Δ = 0.736, *p* < 0.001), mid-luteal vs. early follicular (Δ = 3.534, *p* < 0.001), and mid-luteal vs. peri-ovulatory (Δ = 2.798, *p* < 0.001). The hormonal profile was defined as the first principal component of the levels of E, P, LH, and FSH. The first principal component of hormone levels accounted for 51.37% of the total variance in hormonal fluctuations across sessions. These findings are illustrated in Figure 1.

**Figure 1:**
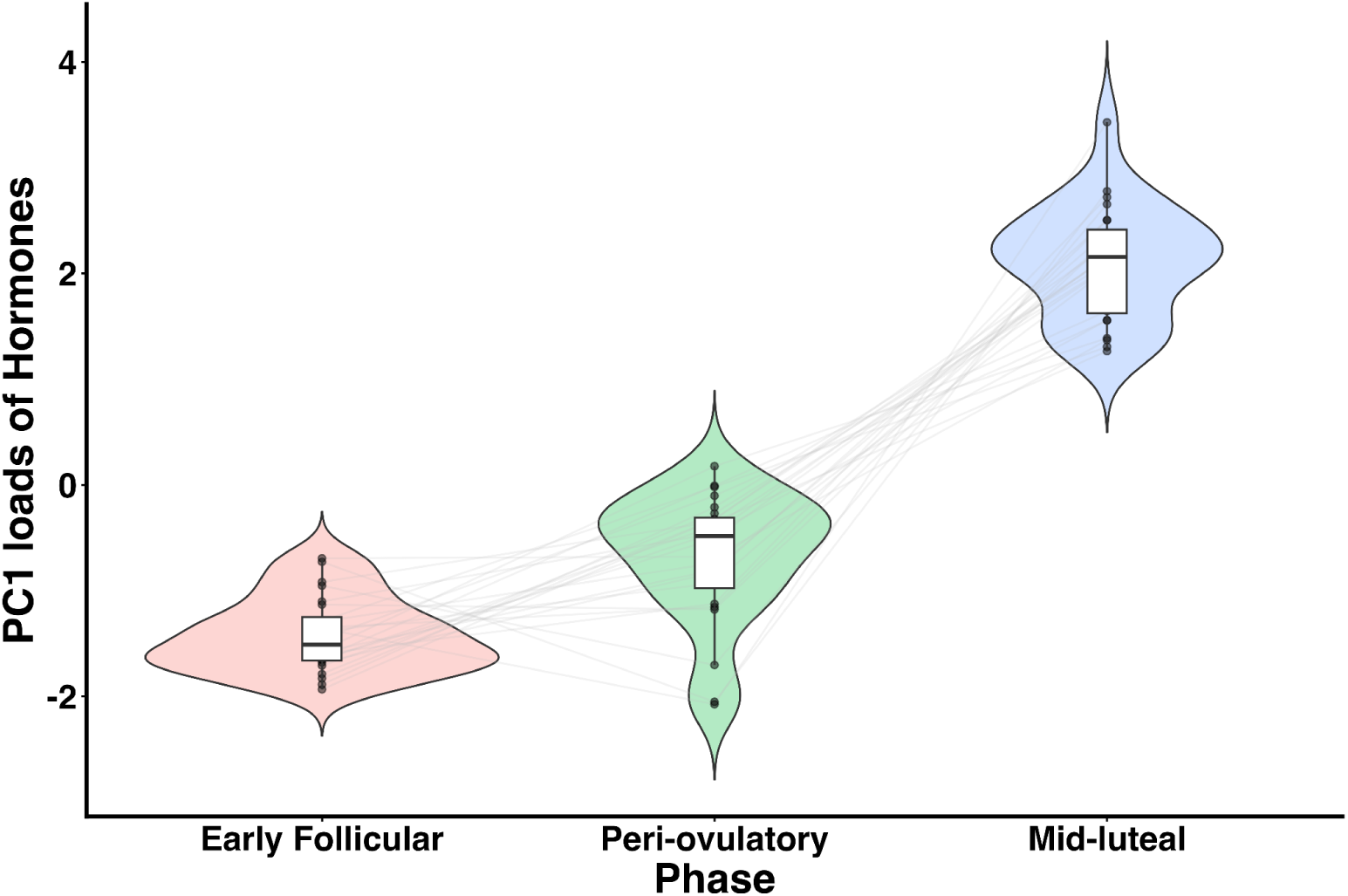
Distribution of the first principal component (PC1) of hormone concentrations across MC phases. The violin plot illustrates the distribution of PC1 values derived from E, P, LH, and FSH. The plot highlights the significant difference in PC1 values across phases, reflecting the typical hormonal fluctuations of the MC. Values from individual subjects are represented by black dots connected through grey lines.

### Microstate analysis

We aimed to identify one or more microstates whose temporal dynamics varied systematically across the three phases of the MC, thereby serving as a potential electrophysiological signature. Microstate analysis was conducted on source-reconstructed resting-state MEG data acquired from 24 participants, each of whom was assessed in all three MC phases. A pooled dataset comprising 72,000 Global Field Power (GFP) peaks was extracted across participants and phases. These GFP topographies were clustered using k-means with the number of clusters (k) ranging from 2 to 40. For each value of k, individual microstate sequences were reconstructed separately for each participant and phase, and the number of visits (i.e., occurrences) to each microstate was quantified. The optimal number of clusters (k = 13) was determined using a data-driven criterion: we selected the solution that yielded the microstate exhibiting the greatest change in visit frequency across the MC phases (see <u>panel A,</u> Figure 2). Within this optimal solution, two microstates (HDM 0 and HDM 1 <u>panels D and E,</u> Figure 2) demonstrated significant phase-dependent differences in their temporal dynamics (F(2,46) = 14.159, p < 0.0001; F(2,46) = 5.50, p = 0.0074, see Figure 3). Crucially, these states were present in every subject and every phase, despite being derived from a clustering performed on the aggregated dataset (Figure 4 and Figure S3 for HDM 1).

**Figure 2:**
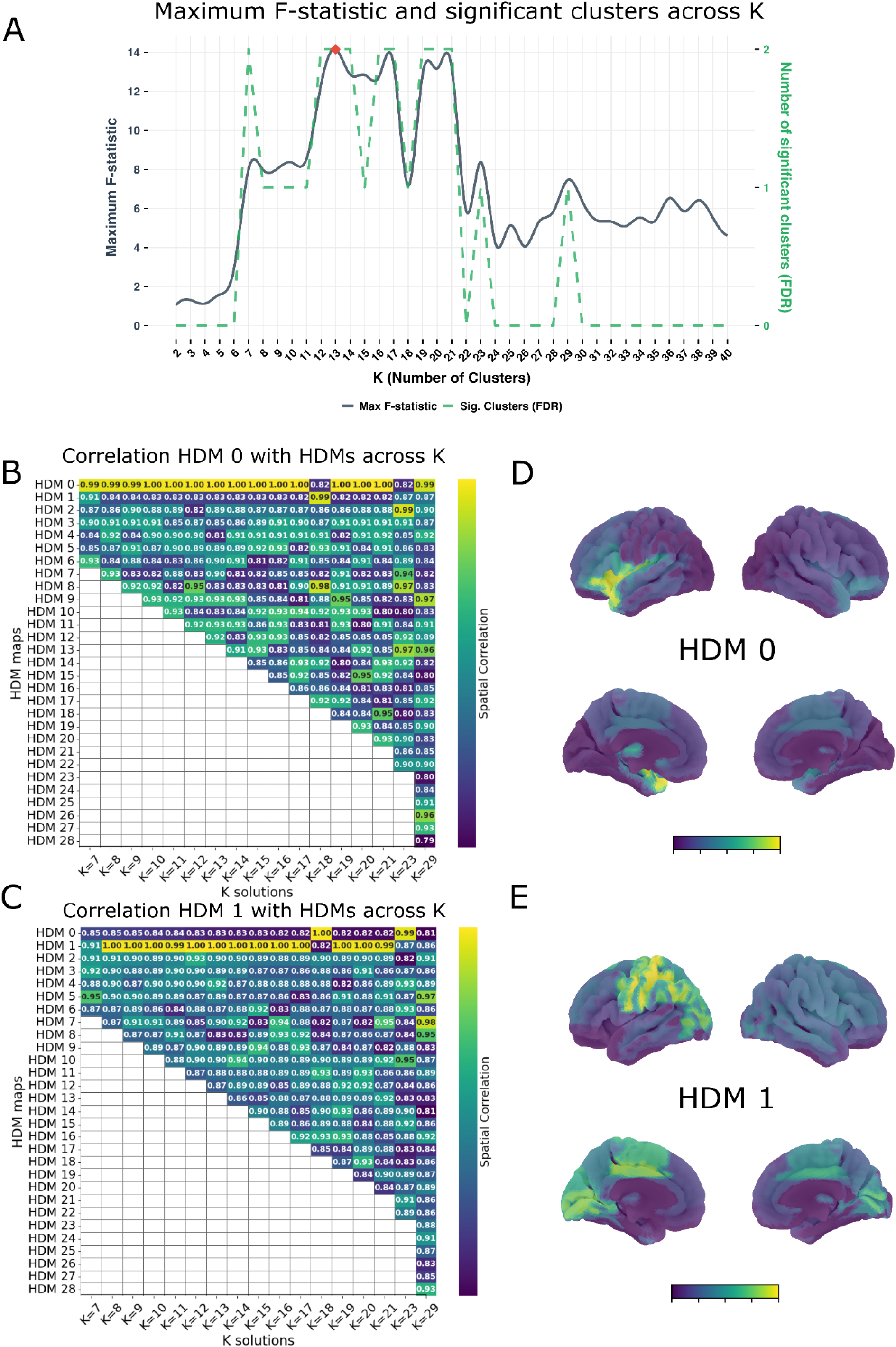
Statistical selection and characterization of Hormone-Dependent Microstates (HDMs). (A) Optimization of the cluster parameter k using the F-statistic criterion (solid black line). The optimal solution was identified at k=13, corresponding to the maximum F-statistic for microstate “visit” frequency across menstrual cycle (MC) phases. The dashed green line indicates the total number of significant F-statistics identified for each value of k following False Discovery Rate (FDR) correction. (B, C) Spatial stability and robustness analysis for HDM 0 and HDM 1. Heatmaps illustrate the spatial correlation between the significant HDMs identified in the optimal k=13 solution and all HDMs across alternative cluster dimensions (k-values) for which at least one significant map exists. The x-axis shows all cluster solutions yielding at least one significant HDM; the y-axis indicates the corresponding number of HDM(s). HDM 0 (B) demonstrated superior robustness, maintaining high spatial correlation across the range of significant k solutions. (D, E) Topographic representations of HDM 0 and HDM 1 for the k=13 solution. The maps display normalized eigenvector weights, with the color scale representing the gradient from minimum to maximum intensity.

**Figure 3:**
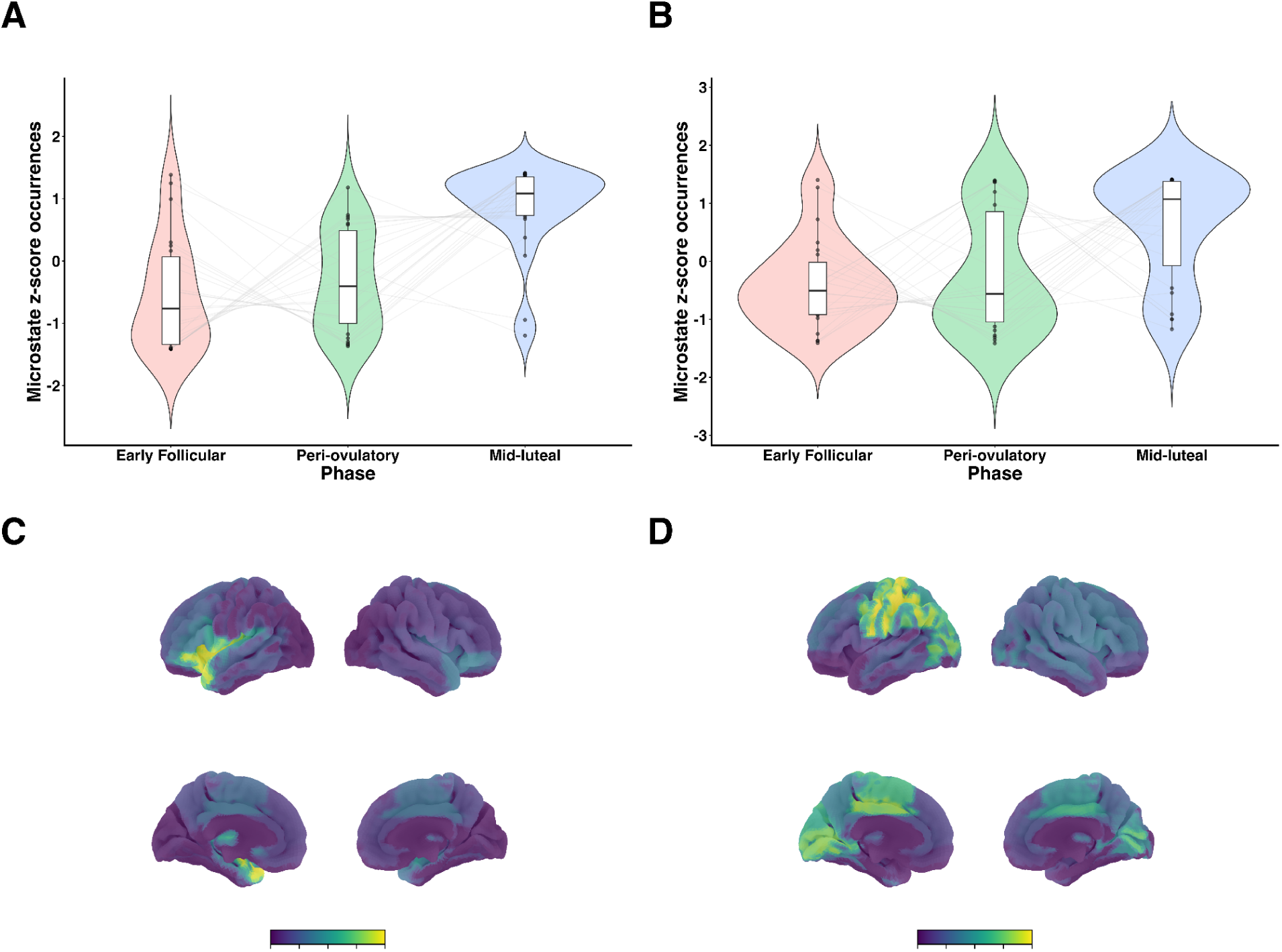
Violin plots illustrating the distribution of occurrences for HDM 0 map (panel A violin plots and C corresponding topographic map) and HDM 1 map (panel B violin plots and D corresponding topographic map) across the three MC phases (early follicular, peri-ovulatory, and mid-luteal). HDM 0 map showed the most pronounced difference in occurrences across the phases. Values from individual subjects are represented by black dots connected through grey lines. The legend of the topographical maps represents the values of the eigenvector weights from minimum to maximum.

**Figure 4:**
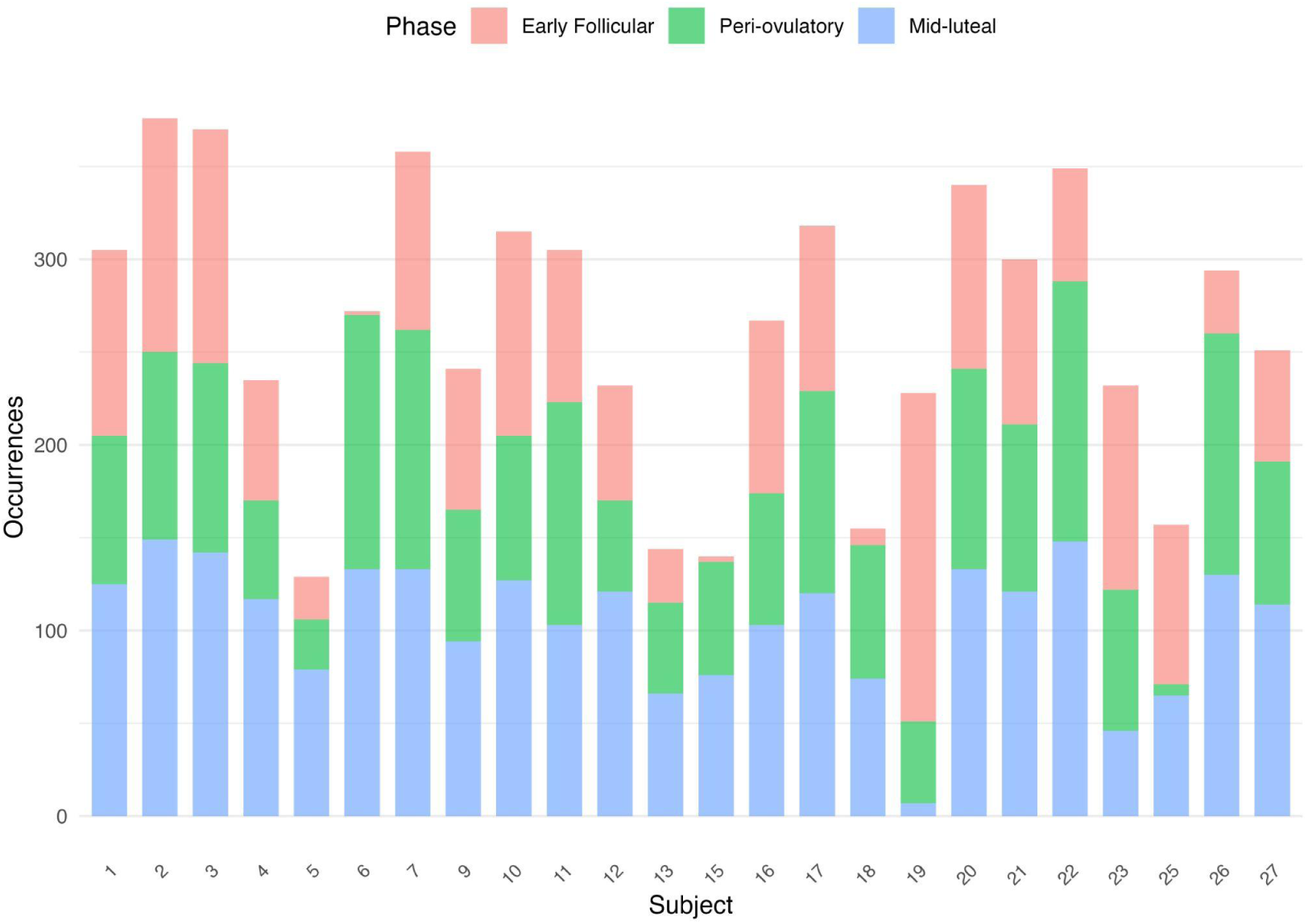
HDM 0 occurrences across subjects and MC phases. The stacked bar plot illustrates the total occurrences of the HDM 0 map for each of the 24 subjects. This visualization confirms that the HDM 0 map is expressed in all participants across all three recorded MC phases, demonstrating its consistent presence.

Post-hoc analyses revealed statistically significant differences in the total number of occurrences of both HDMs (HDM 0 and HDM 1) between the mid-luteal phase and the other two phases (early follicular and peri-ovulatory; see Table 1 and Table 2). HDM 0 was retained for subsequent analyses, given the more robust phase-dependent modulation, that is, the more persistent statistical significance across varying clustering granularities (k’s) (<u>panels B and C</u> Figure 2).

**Table 1:**
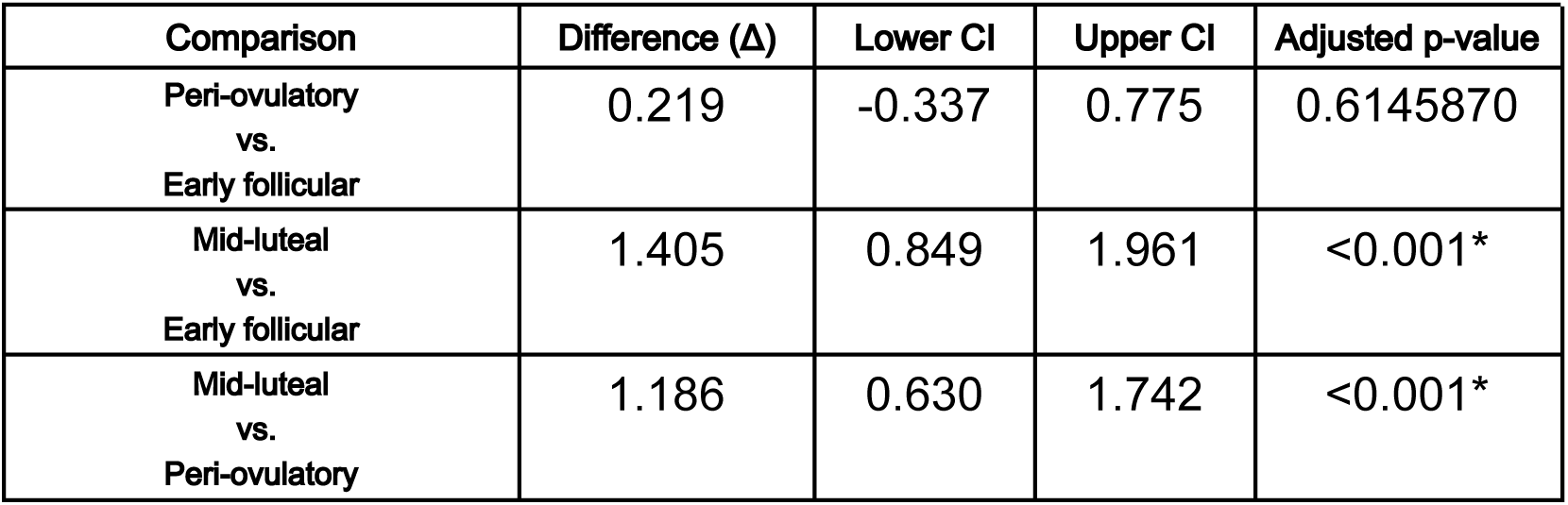
Post-hoc Tukey’s HSD test results for HDM 0 map occurrences across MC. Significant comparisons are marked with p < 0.05.

**Table 2:** Post-hoc Tukey’s HSD test results for HDM 1 map occurrences across MC. Significant comparisons are marked with p < 0.05.

| Comparison | Difference ($\Delta$) | Lower CI | Upper CI | Adjusted p-value |
| --- | --- | --- | --- | --- |
| Peri-ovulatory<br>vs.<br>Early follicular | 0.178 | -0.457 | 0.813 | 0.781 |
| Mid-luteal<br>vs.<br>Early follicular | 1.005 | 0.370 | 1.640 | <0.001* |
| Mid-luteal<br>vs.<br>Peri-ovulatory | 0.827 | 0.192 | 1.462 | <0.001* |

The spatial topography of HDM 0 was predominantly left-lateralized, with its strongest projections localized to the temporal pole, pallidum, insula, amygdala, putamen, inferior frontal orbital gyrus, Rolandic operculum, and thalamus. Similarly, HDM 1 exhibited a left-hemispheric predominance, with peak projections over the precentral and postcentral gyri, inferior parietal gyrus, angular gyrus, cingulate cortex, as well as the lingual and occipital gyri. Globally, all thirteen states (HDMs) for the optimal solution correspond to a specific topography, expressed as a map of regional participations (see Figure 5). The HDM maps corresponding to three of the thirteen HDMs were bilateral, encompassing the frontal and temporal cortices, the precuneus, and the cingulum, respectively. The remaining ten HDM maps exhibited homologous, lateralized networks.

**Figure 5:**
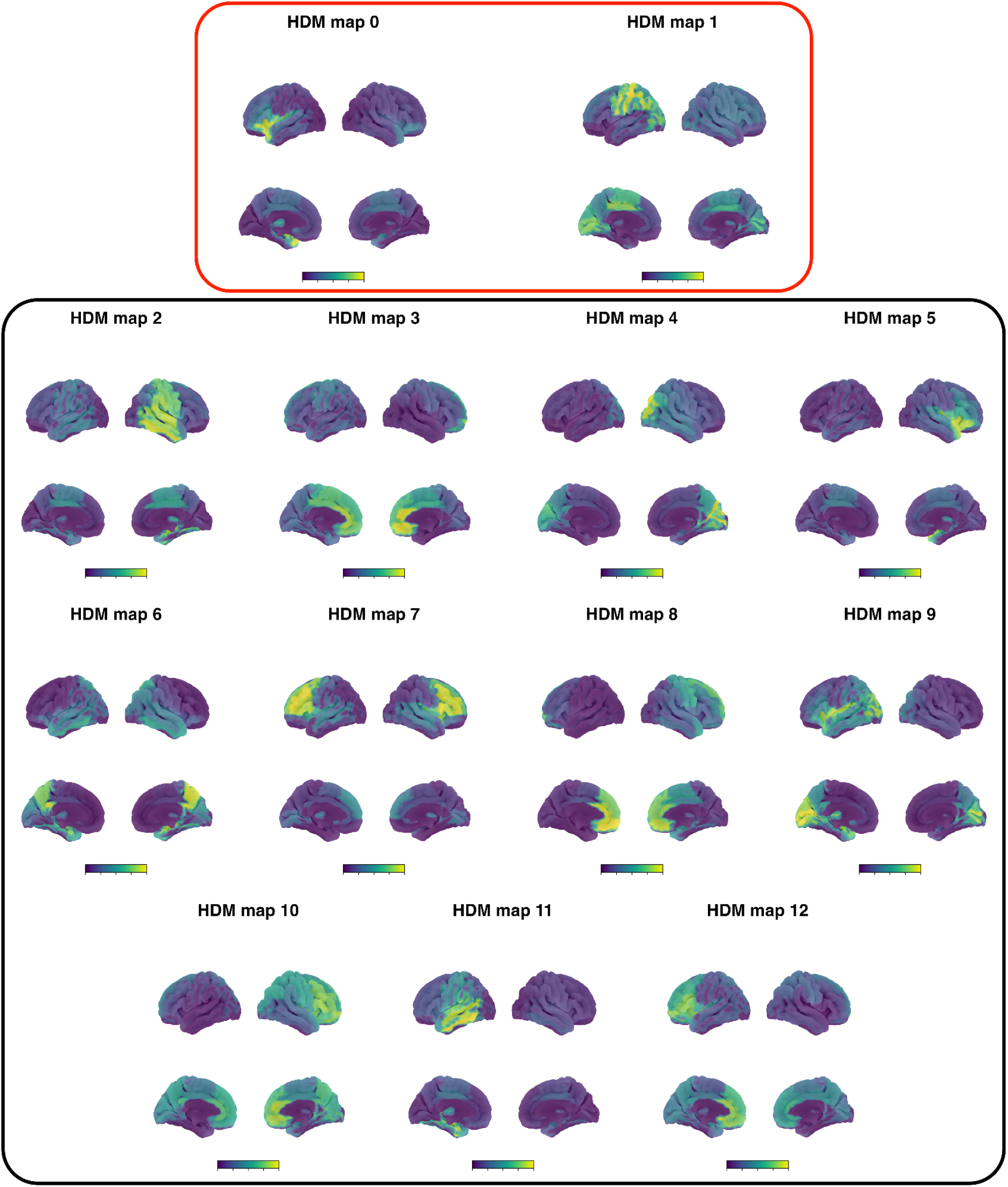
The 13 spatial Hormone-Dependent Microstate (HDM) maps derived from the empirical MEG source-level data and sorted according to F-statistic. The legend represents the values of the eigenvector weights from minimum to maximum. The two significant maps are shown in the red box: HDM 0 and HDM 1. Three out of 13 maps (HDM 7, HDM 6, and HDM 3) are bilateral, involving the frontal cortices, the temporal cortex and precuneus, and the cingulum. The remaining ten maps show homologous, lateralized networks, and are organized into five pairs: (0, 5),(2, 11) (12, 8), (1, 10), and (9, 4).

### Microstates and hormone fluctuation

After demonstrating differences in HDM occurrence across the MC, we now set out to test the hypothesis that these fluctuations are related to hormonal fluctuations. Hormonal fluctuations significantly improved the prediction (based on a linear mixed model - LMM) of the occurrences of microstates compared to an intercept-only model (likelihood ratio: χ^2^ (1) = 40.02, *p* < 0.001). Specifically, hormonal PC1 emerged as a robust positive predictor of HDM 0 occurrences (β = 0.407, 95% *CI* [0.297, 0.518], *t*(70) = 7.21, *p* < 0. 001). The marginal *R*^2^ indicated that hormonal shifts accounted for 42.3% of the variance in microstate activity.

### Microstate, Hormones, and Psychological scores

Hormonal levels and HDM occurrence are related, as demonstrated above. We now set out to test the hypothesis that subject-level mood changes along the MC are predicted by both hormonal levels and microstate dynamics. We compared the null model (*personal growth* ~ *age* + *education* + (1 | *subject*) against a full model incorporating microstate occurrence and hormonal PC1 to predict the variations in the feeling of personal growth (*personal growth* ~ *age* + *education* + *hormonal_PC_*_1_ + *HDM* 0*_visits_* + (1 | *subject*). The full model improved over the null *X*^2^(1) = 7.60, *p_uncorrected_* = 0.02. In particular, hormones and HDM 0 accounted for approximately 7% of the variance for personal growth score (*R*^2^*_marginal_* = 0.07). The AIC for the full model was lower than the baseline (*null*: 463.55 *vs full*: 459. 95, Δ*_AIC_* = 3.60), suggesting improved predictive power. However, when applying the more stringent BIC, which penalizes model complexity more heavily, the null model was slightly preferred (*null*: 474. 94; *full*: 475. 89). Note that, after applying FDR correction across the six Ryff sub-dimensions, this relationship was interpreted as a non-significant trend (*p_FDR_* = 0.13). No other sub-dimensions of well-being showed significant improvement in fit relative to the null model.

To confirm the stability of these estimates, the full model was validated using leave-one out cross-validation (LOOCV) approach. The cross-validation yielded a mean squared error (MSE) of 17.97 and confirmed high internal consistency (average *R*^2^*_conditional_* = 0.868; *R*^2^*_marginal_* = 0.06), as evident by the high degree of overlap between the observed and predicted values in Figure 6.

**Figure 6:**
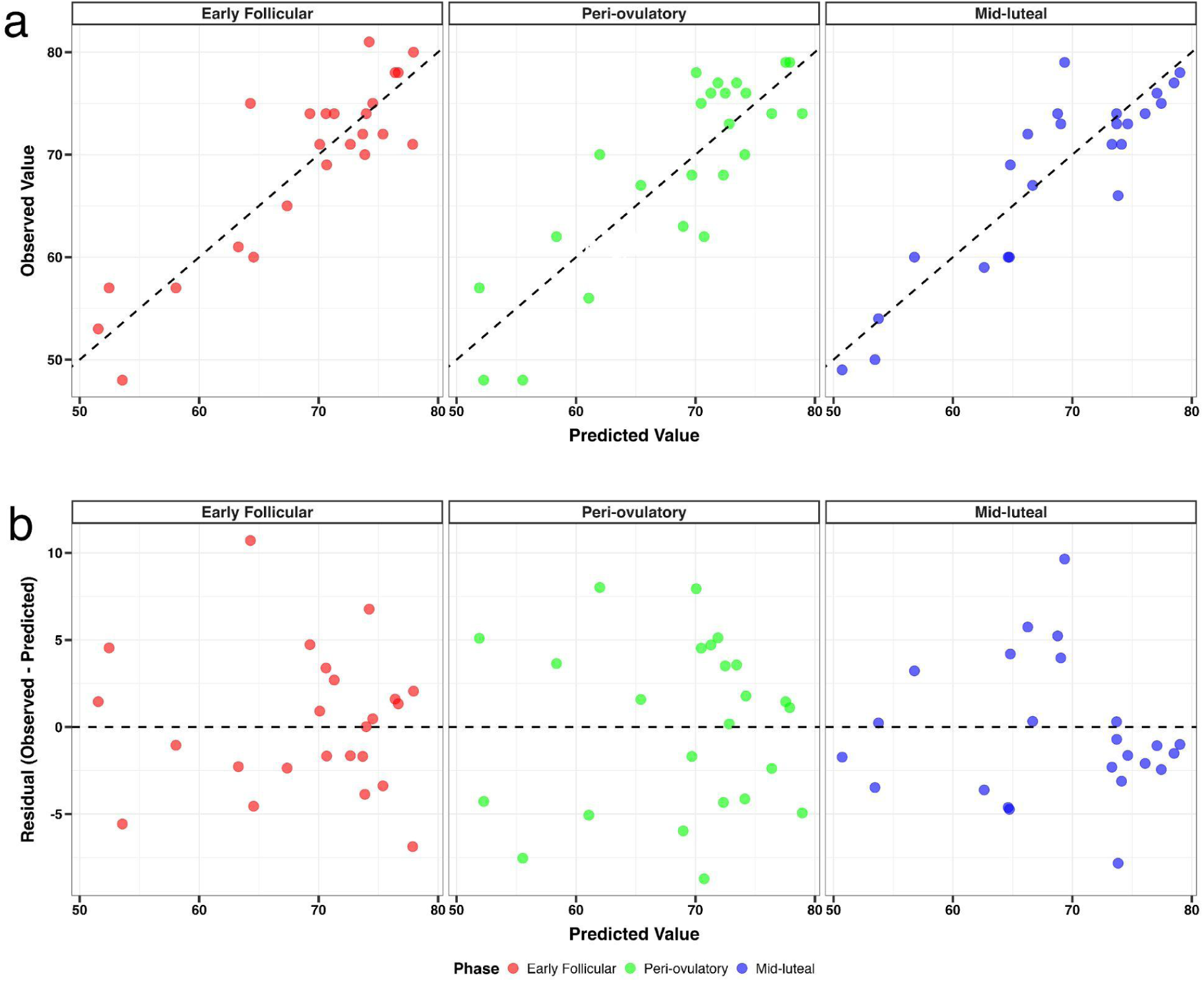
Observed versus predicted personal growth scores. Panel (a) illustrates the distribution of observed and predicted personal growth scores across the three MC phases, derived from the Leave-One-Out Cross-Validation (LOOCV) procedure. Panel (b) shows the model residuals stratified by MC phase.

When taking into account the Ryff’s test as a whole (i.e., the aggregated score over the 6 sub-dimensions), hormones and microstates were not statistically significant predictors χ^2^ (1) = 3.25, *p* = 0.20; *null*: *AIC* =684.08, *BIC* = 695.46; *full*: *AIC* = 684.83, *BIC* = 700.76).

## Discussion

This study investigated the influence of MC phases on brain activation patterns using a source-level microstate analysis framework applied to MEG data. The primary objective was to identify and characterize patterns of brain activity that vary across MC phases and the associated fluctuations in sex hormones.

Our findings identified unique microstates (HDMs) that exhibited phase-dependent variations in the number of visits, with a progressive increase from the early follicular through to the mid-luteal phase. Furthermore, the occurrence of this state was robustly associated with systemic hormonal variance, as operationalized by the first principal component (PC1) of the blood levels of estradiol, progesterone, follicle-stimulating hormone, and luteinizing hormone.

Additionally, we explored the relationship between psychological well-being, hormonal dynamics, and microstate visits. Exploratory analyses revealed that the interplay between sex hormone fluctuations and microstate visits showed a significant association with the personal growth subscale of Ryff’s Psychological well-being scale^35^.

The results align with prior research, such as studies by Liparoti et al.^10,11^, which highlighted the impact of MC phases on brain connectivity and dynamics. While Liparoti et al.^11^ did not establish a direct link between sex hormones and brain dynamics, in the current study we identified a microstate that provides a brain signature associated with hormonal fluctuations, as reflected in the first principal component of hormone levels. Specifically, whereas Liparoti et al. reported changes in the overall flexibility of brain dynamics, the present analysis identifies specific topographies that relate to hormonal levels and psychological states.

While both studies aimed to characterize the brain’s dynamic state exploration across the MC, they employed distinct frameworks. Microstates, defined as “quasi-stable” periods of consistent electrical topography across MEG source-level time-series ^32^, provide a global perspective by encompassing the entire brain. In contrast, neuronal avalanches, as described by Liparoti et al., represent cascades of neural events originating from a single brain region and propagating through the network^36–38^. These methodological distinctions likely capture different facets of the underlying neurophysiological processes, offering complementary insights into the interplay between hormonal fluctuations and large-scale brain activities. This aligns with findings from De Filippi et al.^39^, who reported phase-dependent differences in information processing related to fluctuations in estradiol and progesterone, specifically between the luteal and follicular phases. Collectively, these diverse frameworks suggest that hormonal shifts do not merely alter localized activities but reorganize the brain dynamics at the large scale.

The observed patterns are consistent with earlier reports, such as those by Becker and Bazanova^40,41^, which documented a frequency shift in the alpha power spectrum between the follicular and luteal phases, with faster rhythms detected in the latter. These spectral changes, likely influenced by progesterone fluctuations^41,42^, may suggest a mechanistic link to the current results, as systematic variations in local EEG spectral amplitude have been associated with microstate occurrences and dynamics^43^.

Furthermore, the findings extend the work of Cacioppo et al. ^33^, who identified MC-dependent hemispheric asymmetry using microstate analysis. Their study suggested that specific microstates, such as those with prominent Left Anterior–Right Posterior topographies, modulate responses to emotional word presentation during the menstruation phase as compared to the early luteal phase.

The convergence of these data underscores a sophisticated interplay among endocrine fluctuations, neural dynamics, and the multifaceted dimensions of human behavior^42,44,45^. Our findings—magnetoencephalographic MC marker (i.e., the frequency of visits to specific global configurations) and hormonal variance together predicting subtle shifts in ‘personal growth’—support a tripartite framework in which neurophysiological states act as a critical mediator between the hormonal milieu and psychological well-being. While our study is specifically concerned with the healthy population, our results might also be relevant for clinical populations, such as those with Premenstrual Dysphoric Disorder (PMDD), where a heightened sensitivity to hormonal fluctuations could manifest as more robust alterations in both microstate visits and their corresponding behavioral outcomes^1,4^.

Methodologically, the present study builds upon the analytical framework established by Tait et al. (2022)^32^, incorporating several notable modifications.

First, our approach to determining the optimal number of clusters deviated from standard microstate analysis, which often employs subjective measures such as the k-needle algorithm or similar criteria^32,34^. Instead, we tailored our selection of the optimal cluster number based on the MC phases. This allowed us to select microstate maps (HDMs) that maximize contrast across these phases, thereby highlighting the activation patterns most relevant to our hypothesis. Crucially, we demonstrated that these specific microstate maps were ubiquitous across all subjects. Furthermore, they exhibited anatomical properties, such as lateralization and homology, consistent with findings from other MEG source-level microstate studies^32,46^.

Second, we did not include traditional microstate metrics in our analyses, such as mean microstate duration, percentage of time covered, or transition probabilities, to characterize microstate dynamics. This decision was informed by the aim to prioritize simpler features^47^, thereby minimizing potential biases introduced by arbitrary methodological choices (e.g., assigning data points proximal to GFP peaks to specific microstate classes^48^) and mitigating the risk of feature intercorrelation^49^.

The present study has several limitations. While this is the largest MC longitudinal MEG dataset to date, the sample size might not allow to capture effects with small sizes. This might be evident in particular for the results of the predictive models, given that in fact, the association identified was no longer significant after correction for multiple comparisons. Along these lines, the Ryff’s scale may lack the sensitivity required to detect subtle differences across the MC in our sample (see Figure S1 and Figure S2 in supplementary materials), which consisted of women without mood disorders (such as PMDD). It is worth noting that even when mood disorders are present, psychological changes tend to be inconsistent and fragmented^4^. Given that personality traits are mostly stable across the MC (e.g., being optimistic), a small sample size might miss subtle changes, as most of the variance, as expected, is captured by the random effect of the subject.

Second, sampling the MC at three discrete time points limits our ability to sample individual variability across multiple menstrual cycles, which would require tracking the participants over multiple months. However, this dataset remains the only available longitudinal MEG dataset across the MC. Third, the study exclusively focused on naturally cycling women, thereby limiting the applicability of the findings to other populations, such as individuals using hormonal contraceptives or those with irregular menstrual cycles.

Notwithstanding these limitations, the study provides novel insights into the neurophysiological mechanisms underlying MC-related changes in brain function. The findings underscore the pivotal role of sex hormones in modulating large-scale neural dynamics and their potential influence on behavior. We propose that the dynamics of the brain microstates may be mediated by hormonal fluctuations associated with the MC, and might be seen as neurophysiological and dynamical substrates of subtle psychological changes occurring over the MC. Lastly, these results contribute to the growing body of literature emphasizing the importance of accounting for hormonal fluctuations as a yet-unaccounted-for source of variability in any neuroimaging study that includes women in fertile age^50^.

Finally, our results talk to the idea that the brain should not be treated in isolation with respect to other bodily systems, and shows how ultra-slow dynamics, in this case at the month time-scale, contribute and constrain significantly faster (i.e., millisecond) dynamics dynamics.

## Methods

### Participants

We used a subset of subjects as described in (Liparoti et al. 2024, 2021): 24 right-handed, heterosexual, native Italian-speaking females with regular MCs, aged (26.36 ± 5.07) years, and with (16.52 ± 2) years of education. Participants provided informed consent, and the study was approved by the Local Ethics Committee of the University of Naples “Federico II” (protocol n. 223/20). Women were excluded if they had a history of neuropsychiatric disorders, premenstrual dysphoric symptoms, recent pregnancy, or hormonal contraceptive use in the six months prior to the study. To minimize confounding factors, participants abstained from tobacco, alcohol, and caffeine for 48 hours before MEG recordings, which were conducted at consistent times to control for circadian influences.

### Experimental protocol

Participants were evaluated during three distinct phases of the MC, as outlined in Liparoti et al. 2024, 2021^10,11^: the early follicular phase (cycle days 1–4, characterized by low levels of estradiol and progesterone, session 1), the peri-ovulatory phase (cycle days 13–15, marked by elevated estradiol levels, session 2), and the mid-luteal phase (cycle days 21–23, characterized by high levels of both estradiol and progesterone, session 3). The timing of these phases was determined using the back-counting method, with the self-reported onset of menses serving as the reference point to estimate the peri-ovulatory and mid-luteal windows. During each MC phase, MEG recordings and blood samples were collected to measure concentrations of sex hormones, including E, P, FSH, and LH. Additionally, a transvaginal pelvic ultrasonography examination was conducted during the early follicular phase to verify the correctness of the time point, and structural magnetic resonance imaging (MRI) was performed following the final MEG session. To control for potential recording session effects, the order of the MC phases across the three recording sessions was randomized. Detailed methodologies for hormone analysis, pelvic ultrasonography, and MRI acquisition are provided in the supplementary materials of Lipatori et al. (2024, 2021)^10,11^. Hormonal concentration data were available for 24 of the 26 participants.

### MEG recordings

MEG data were acquired using a 163-channel system (154 magnetometers and 9 reference channels) based on superconducting quantum interference devices (SQUIDs) and housed within a magnetically shielded room (AtB Biomag UG, Ulm, Germany), as detailed in Liparoti et al.^10,11^. This system is characterized by a magnetic field noise spectral density of approximately 5 fT/Hz 1/2. Prior to data acquisition, the participants’ head positions were digitized using four anatomical landmarks and four position coils to ensure accurate co-registration. Each participant underwent two eyes-closed MEG recordings per phase (corresponding to early-follicular, peri-ovulatory, and mid-luteal phases), each lasting 3 minutes and 30 seconds (sample frequency 1024Hz), with a brief inter-session rest period during which participants remained seated within the MEG room. Electrocardiogram (ECG) and electro-oculogram (EOG) signals were simultaneously recorded to facilitate the identification and removal of physiological artifacts.

For subsequent analyses, only the first 3 minutes and 30 seconds of MEG data were utilized. These data were preprocessed using automated pipelines as described in Liparoti (2021, 2024)^10,11^. Preprocessing steps included principal component analysis (PCA) to attenuate environmental noise, implemented in MATLAB using the FieldTrip toolbox, and independent component analysis (ICA) to remove physiological artifacts, such as cardiac and ocular signals. Source reconstruction was performed using the Linearly Constrained Minimum Variance (LCMV) beamformer approach ^51^, employing the volume conduction model proposed by Nolte et al. (2003) ^52^. The Automated Anatomical Labeling (AAL) atlas was used to define 90 cortical regions of interest (ROIs), excluding cerebellar regions due to potential concerns about signal reliability. Beamformed time-series data were visually inspected, and only artifact-free segments were retained for further analysis. Segments shorter than 2 seconds were excluded.

### A Source-Space Microstate analysis

The analytical pipeline employed in this study builds upon the source-space microstate analysis framework proposed by Tait et al.^32^, with several methodological adaptations. Notably, the source reconstruction in our study utilized the Linearly Constrained Minimum Variance (LCMV) beamformer approach, whereas Tait et al. employed the eLORETA method. Additionally, the parcellation scheme adopted in this work was based on the Automated Anatomical Labeling (AAL) atlas, encompassing 90 regions of interest (ROIs), in contrast to the HCP230 atlas used in Tait et al.’s analysis. Furthermore, the source-reconstructed data in our study were band-pass filtered between 2–30 Hz for subsequent microstate analysis.

### Extraction of GFP Peaks

To improve the signal-to-noise ratio and ensure the topographic consistency necessary for clustering, we identified and selected time points corresponding to peaks in the Global Field Power (GFP). GFP was defined as the standard deviation of the signal across ROIs. Prior to GFP computation, the beamformed time-series for each participant were standardized using z-scores for each phase. Additionally, to enhance the robustness of peak detection, the GFP signal was smoothed using a moving average filter with a window length of five samples. Finally, to mitigate the effects of potential source orientation flipping across subjects^32^, the absolute values of the z-scored source estimates were utilized following peak extraction.

For group-level analysis, the *1,000 GFP peaks* with the highest heights were sampled from each participant and each phase and pooled together. GFP peaks that were apart less than 10 samples (~0.01s) were discarded.

Given the inclusion of 24 participants and three phases per participant, a total of *72,000 GFP peaks* were aggregated and subsequently submitted for k-means clustering using the Python analysis toolbox Neurokit2^53^.

### k-means Clustering

The modified k-means clustering algorithm^54^ was executed ten times with random initializations, and the iteration yielding the highest Global Explained Variance (GEV) was selected. Following this, eigenvector decomposition was used to refine the cluster centroids^32,54^.

In traditional microstate analysis, the optimal number of clusters is selected by comparing cluster performance metrics, such as GEV or cluster separability, across different numbers of microstates. Using this approach allows to describe global brain activity through a series of generalizable “building blocks” or activity patterns. However, this study diverged from “traditional” microstate analysis^32,34^ and, rather than describing global brain activity patterns, aims to identify specific microstates (HDMs) that vary as a signature of the human MC.

To this end, an alternative approach was employed to select the optimal number of clusters (k) by identifying the microstate maps that best distinguished between the different MC phases. Specifically, for each chosen k (number of clusters) and for each of the k microstate maps, an F-statistic was calculated to determine if the number of occurrences (“visits”) to a particular states differed across the phases of the menstrual cycle. The partition that yielded the state with the highest F-statistics was then selected.

F-statistics were computed using a repeated measures analysis of variance (ANOVA). Note that, for each choice of k clusters, the ANOVA was performed k times (one per microstate). To account for the multiple comparisons, p-values were adjusted using the False Discovery Rate (FDR) correction method^55^. In cases where the ANOVA revealed statistically significant results, post-hoc pairwise comparisons between MC phases were performed using Tukey’s Honest Significant Difference (HSD) test to control for multiple comparisons. The statistical analysis for the repeated measure ANOVA was performed using the Python package statsmodels^56^.

By employing this strategy, we isolated microstate maps whose temporal dynamics—specifically the number of visits to specific global configurations—varied significantly as a function of the MC phase (HDMs). The map providing the most statistically significant differentiation between phases was selected for further analysis. It is noteworthy that, although the HDMs were defined using a grand-concatenated dataset, the phase-sensitive map was identified in every participant at each recording time point.

### Psychological evaluation

Psychological assessments were conducted during each MC phase. Detailed descriptions of the assessment procedures can be found in Liparoti et al.(2021)^10^. For the purposes of this study, we utilized data from the cumulative Ryff’s Psychological Well-Being Scale, which evaluates six dimensions of well-being: autonomy, environmental mastery, personal growth, positive relations with others, purpose in life, and self-acceptance^35^.

### Statistical Analysis

#### Association between microstate occurrences and hormonal concentrations

To examine the relation between hormonal changes and the HDM map, we implemented a Linear Mixed-Effects Model (LMM). The LMM utilized the z-score of occurrences (for each subject across phases) as the dependent variable. The first principal component of hormones (PC1) served as the fixed predictor, with ‘subject’ included as a random effect. In more detail, the PCA was conducted on the hormonal concentration variables (E, P, LH, and FSH) to reduce dimensionality and mitigate multicollinearity among the sex hormones. Subject-level variability was first removed to ensure independence of observations for the PCA, given the repeated-measures design.

#### Association between microstate occurrences, hormonal concentration, and psychological evaluation

To assess the interplay between psychological well-being, HDM occurrences, and hormonal levels, an LMM model was employed. The model accounted for the study’s repeated-measures design, in which 24 participants were assessed across three distinct MC phases. The LLM framework facilitated the inclusion of both fixed effects (predictors of interest) and random effects (to account for inter-individual variability).

To estimate well-being scores, we constructed a null LLM. This model incorporated age and education as time-invariant fixed-effect covariates, with the subject included as a random predictor, and no other fixed predictors. This baseline model was then compared with a full model. This latter included the PC1 of hormones and the z-score of microstate map occurrences as additional fixed predictors. All analyses were performed using LMM in R via the lme4 and lmerTest packages.

Model comparisons between a null and a full model were performed using Likelihood Ratio Tests (LRT). Final parameter estimates were derived using Restricted Maximum Likelihood (REML) with Satterthwaite’s approximation for degrees of freedom. Model quality was further assessed using the Akaike InformationCriterion (AIC), Bayesian Information Criterion (BIC), and Nakagawa’s *R*^2^ (partitioned into marginal and conditional components).

Additionally, a more detailed analysis was incorporated to provide separate estimates for each well-being subscale: personal growth, autonomy, environmental mastery, positive relations, purpose in life, and self-acceptance.

To control the False Discovery Rate (FDR) while maintaining statistical power, p-values were corrected using the Benjamini-Hochberg procedure across the six specific subscales. The global aggregate score (well-being score) was treated as a separate category and was not included in the subscale correction pool.

To assess the generalizability and robustness of our findings, we implemented a Leave-One-Out Cross-Validation (LOOCV) procedure for the well-being domains that could be predicted. For each iteration, the model was trained on N-1 observations and used to predict the value of the omitted observation. Predictive accuracy was quantified using Mean Squared Error (MSE) and the consistency of *R*^2^ estimates across folds.

All the models were trained and assessed in R with the following packages lmerTest, dplyr, performance, MuMIn, ggplot2^57–62^.

All the code for the analysis is available at https://github.com/suforraxi/meg_microstates_menstrual_cycle

https://github.com/suforraxi/microstate_based_on_stats

## Supplementary material: Figures

### Psychological scores

The psychological assessment consisted of Ryff’s test of well-being, which comprises six sub-dimensions (autonomy, environmental mastery, personal growth, positive relations with others, purpose in life, and self-acceptance). While these are stable constructs at the yearly timescale, we aimed to measure small within-subject variations occurring along the MC. However, psychological well-being did not differ significantly across the three MC phases, either globally (Figure S1) (*F*(2, 46) = 0. 213, *p* = 0. 809) or within specific sub-dimensions (Figure S2).

**Figure S1:**
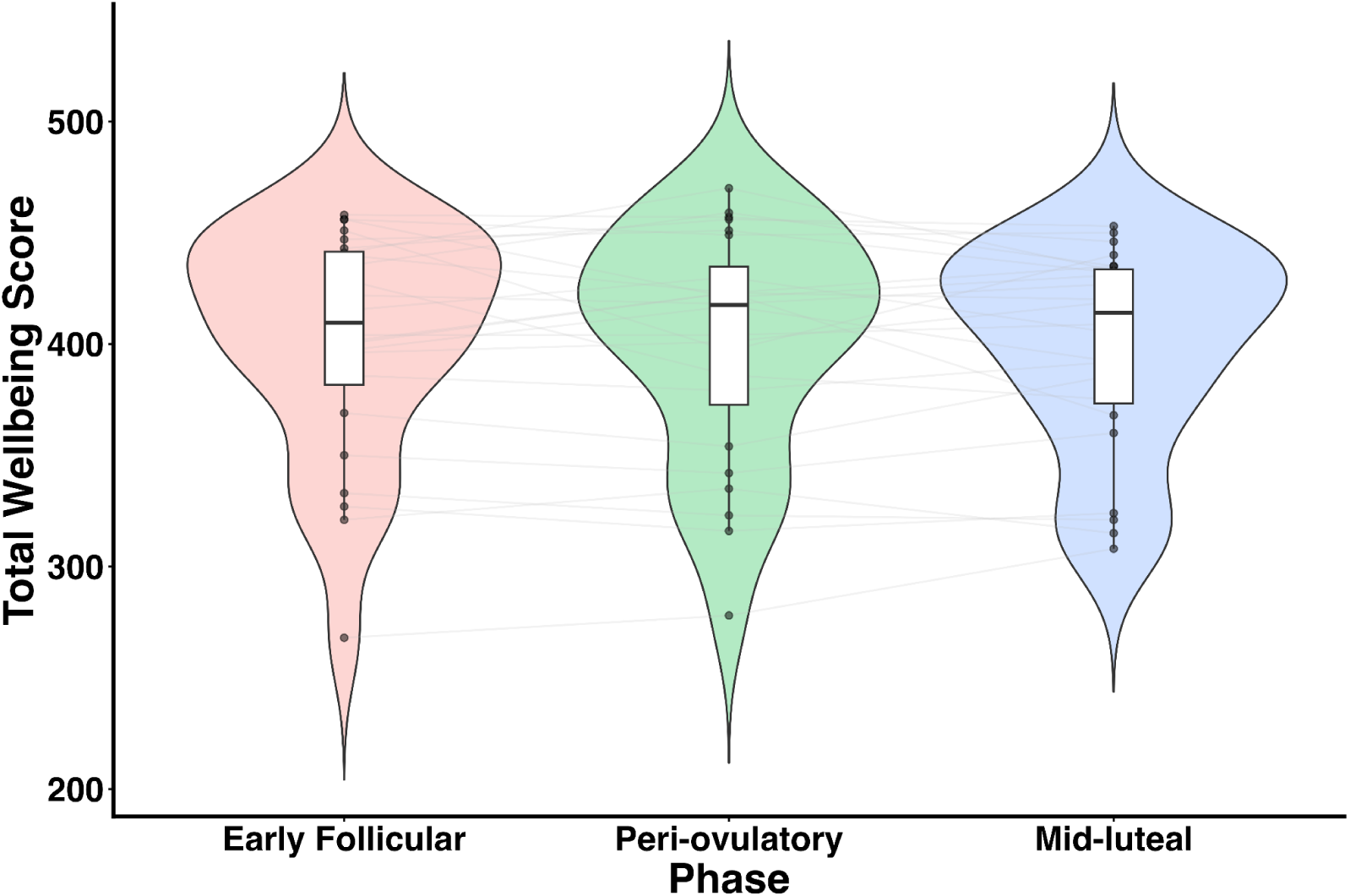
Cumulative Ryff’s test of the six dimensions of well-being.Violin plots show the distribution of the aggregate well-being scores across the three MC phases. The analysis of variance did not reveal a significant effect of MC phase on the cumulative well-being score. Values from individual subjects are represented by black dots connected through grey lines.

**Figure S2:**
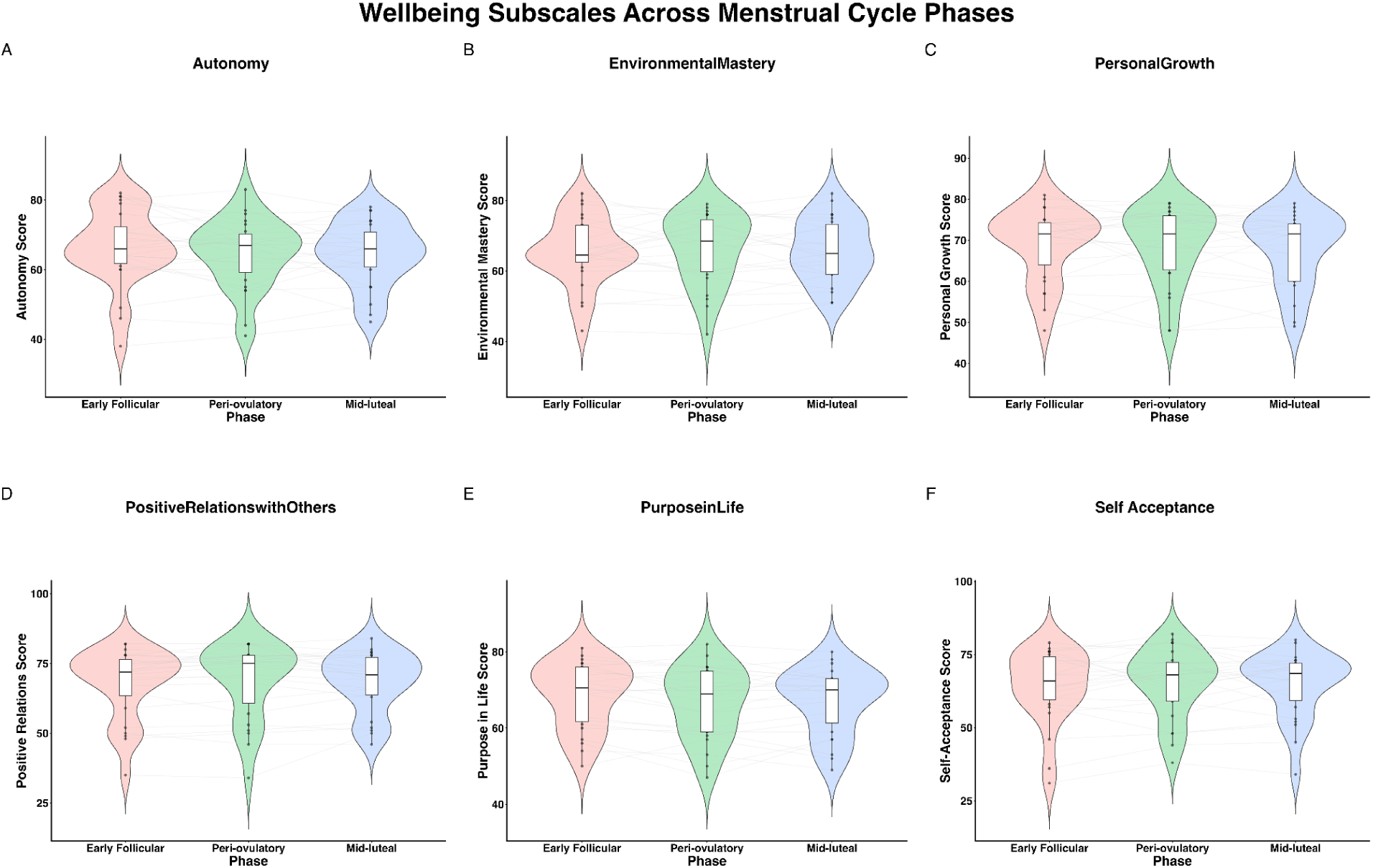
Distribution of the six subscales of Ryff’s Psychological Well-Being. Violin plots illustrate the scores for Autonomy (A), Environmental Mastery (B), Personal Growth (C), Positive Relations with Others (D), Purpose in Life (E), and Self-Acceptance (F) across the three menstrual cycle phases (Early Follicular, Peri-ovulatory, and Mid-luteal). Repeated measures ANOVA revealed no significant effect of the menstrual cycle phase on any of the six individual subscale scores. Values from individual subjects are represented by black dots connected through grey lines.

**Figure S3:**
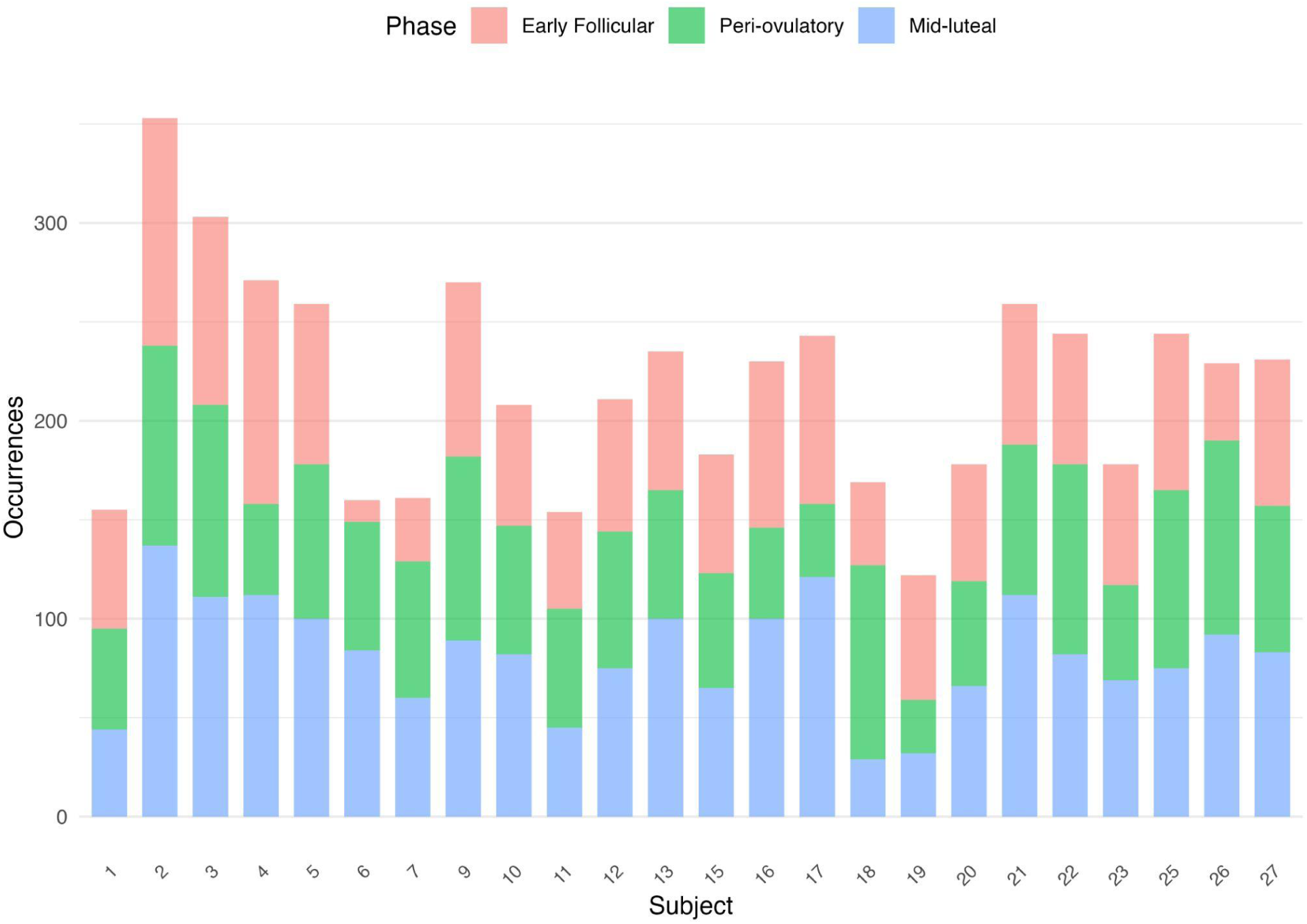
Hormone-Dependent Microstates 1 (HDM) occurrences across subjects and MC phases. The stacked bar plot illustrates the total occurrences of HDM 1 for each of the 24 subjects. This visualization confirms that HDM 1 is expressed in all participants across all three recorded MC phases, demonstrating its consistent presence.

## Supplementary material: Tables

**Table S1:**
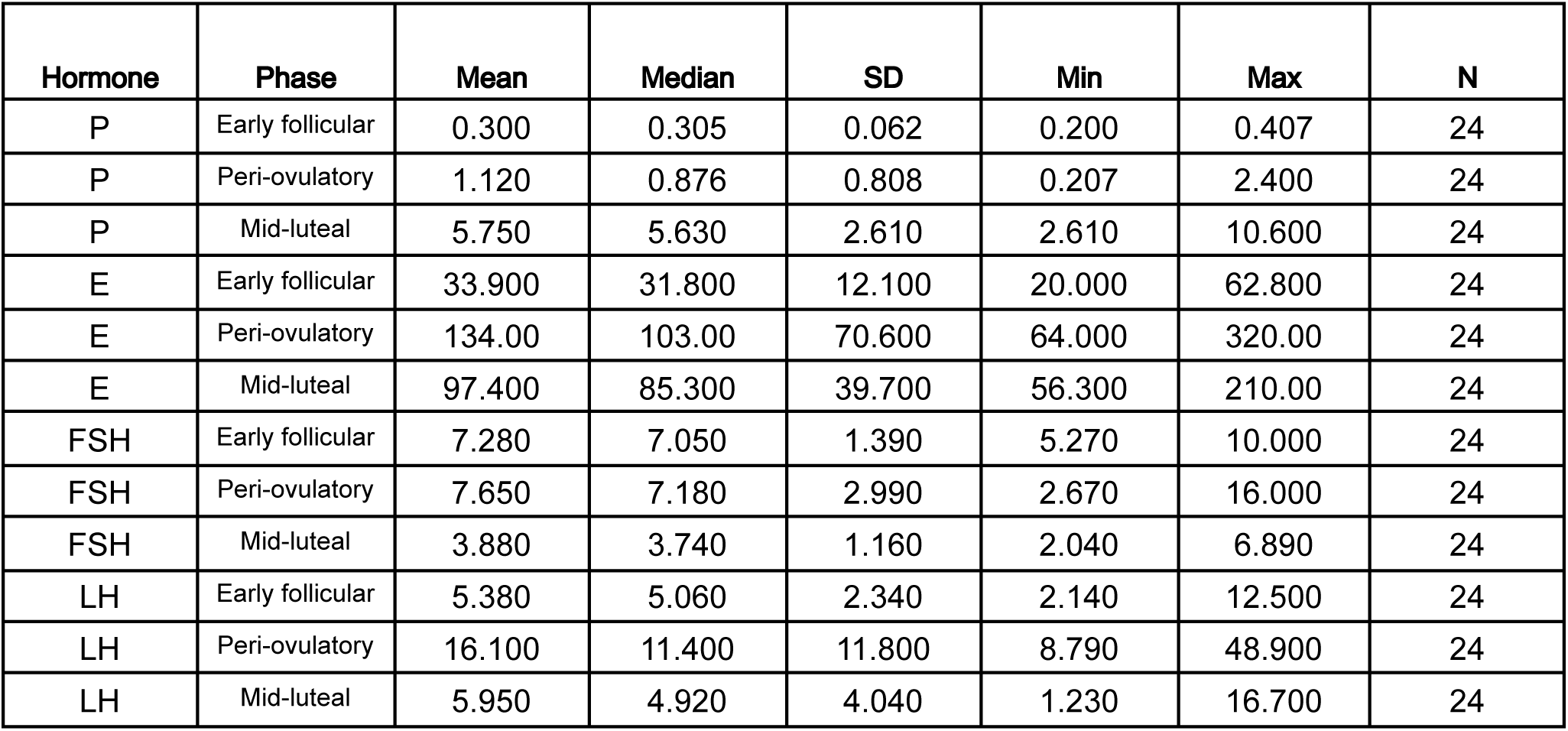
Descriptive Statistics for Sex Hormone Concentrations across Menstrual Cycle Phases. The table presents the mean, median, standard deviation (SD), minimum (Min), and maximum (Max) concentrations for Progesterone (P), Estradiol (E), Follicle-Stimulating Hormone (FSH), and Luteinizing Hormone (LH) during the Early Follicular, Peri-ovulatory, and Mid-luteal phases (N=24).

| Hormone | Phase | Mean | Median | SD | Min | Max | N |
| --- | --- | --- | --- | --- | --- | --- | --- |
| P | Early follicular | 0.300 | 0.305 | 0.062 | 0.200 | 0.407 | 24 |
| P | Peri-ovulatory | 1.120 | 0.876 | 0.808 | 0.207 | 2.400 | 24 |
| P | Mid-luteal | 5.750 | 5.630 | 2.610 | 2.610 | 10.600 | 24 |
| E | Early follicular | 33.900 | 31.800 | 12.100 | 20.000 | 62.800 | 24 |
| E | Peri-ovulatory | 134.00 | 103.00 | 70.600 | 64.000 | 320.00 | 24 |
| E | Mid-luteal | 97.400 | 85.300 | 39.700 | 56.300 | 210.00 | 24 |
| FSH | Early follicular | 7.280 | 7.050 | 1.390 | 5.270 | 10.000 | 24 |
| FSH | Peri-ovulatory | 7.650 | 7.180 | 2.990 | 2.670 | 16.000 | 24 |
| FSH | Mid-luteal | 3.880 | 3.740 | 1.160 | 2.040 | 6.890 | 24 |
| LH | Early follicular | 5.380 | 5.060 | 2.340 | 2.140 | 12.500 | 24 |
| LH | Peri-ovulatory | 16.100 | 11.400 | 11.800 | 8.790 | 48.900 | 24 |
| LH | Mid-luteal | 5.950 | 4.920 | 4.040 | 1.230 | 16.700 | 24 |

